# RNA-sequencing as an expression-informed alternative for somatic variant detection and cancer driver identification

**DOI:** 10.1101/2022.05.10.491431

**Authors:** Jon Akutagawa, Allysia J Mak, Tanvi Damle, Colette Felton, Julie L Aspden, Angela N Brooks

## Abstract

Detecting somatic mutations is a cornerstone of cancer genomics and clinical genotyping, traditionally relying on whole exome (WES) or whole genome sequencing (WGS) data. While RNA sequencing (RNA-seq) is primarily used to assess gene expression, it can also be utilized for somatic variant detection. RNA-seq inherently captures expressed variants, thereby reducing the identification of passenger mutations and mitigating the annotation biases associated with WES. We performed somatic variant calling and cancer driver detection from tumor RNA-seq data from 1,359 RNA-seq samples with matched WGS from the Pan-Cancer Analysis of Whole Genomes (PCAWG) Consortium of the International Cancer Genome Consortium (ICGC) and The Cancer Genome Atlas (TCGA). Our approach using RNA-seq data, alone, identified at least one putative driver gene in 15 of 16 cancer types, consistent with driver predictions from WGS. We further observed that the variant allele frequency (VAF) of RNA-seq–derived variants reflects their predicted functional impact, providing information specific to transcriptomic data. Because RNA-seq data can be generated and analyzed at lower computational cost and time than WGS or WES, it represents a resource-efficient alternative for cancer driver detection. This study provides an extensive pan-cancer resource of RNA-seq-derived variants from the PCAWG consortium and supports the utility of RNA-seq as a complementary approach for characterizing expressed somatic variation in cancer.

## Introduction

The detection of somatic variants through next-generation sequencing (NGS) has enabled researchers and clinicians to associate genetic mutations and disease phenotypes. The rapid improvement and falling costs of these technologies have led to the discovery of a whole host of crucial cancer-driving mutations and have opened new doors for targeted therapies in many cancers. Discovering *EGFR* mutations in lung cancer^1^ and *BRAF* mutations in melanoma^2^ have led directly to novel treatments^3,4^ that have redefined standard of care options for eligible cancer patients. Continued advances in NGS technology, particularly whole-exome sequencing (WES), whole-genome sequencing (WGS), and RNA sequencing (RNA-seq), have allowed researchers to generate massive amounts of NGS data and better understand the genetic basis of cancer. RNA-seq is commonly employed for gene expression and alternative splicing analyses, which has given researchers an opportunity to uncover the transcriptional and post-transcriptional phenotypes of cancer cells.

Existing variant callers have demonstrated the ability to detect novel somatic variants^5^ and can differentiate these somatic mutations from inherited or *de novo* germline mutations, neutral polymorphisms, and artifacts derived from misalignments, sequencing errors, or PCR errors^6–9^. However, these tools are overwhelmingly designed for WES or WGS data. WES employs probes designed for exonic regions, which both introduces annotation biases^10^ and results in the omission of variants within UTRs. Variant detection using RNA-seq data includes such regions of transcripts and therefore represents a more comprehensive search space for variants. Variants found in these transcripts are known to be expressed and are more likely to have a functional impact on the cell phenotype. Despite these advantages, the transcriptome’s inherent complexity can prove to be technically challenging when detecting variants. RNA-seq data often contain reads that span intronic regions or harbor variants in genes with low expression, which pose problems for many of these current variant calling tools^11^. The identification of somatic variants in RNA-seq data has previously been achieved^12–16^, but there are few studies with results validated by a matched whole-genome variant list^17^.

To address this gap, we implemented and evaluated an integrated pipeline for detecting somatic variants in RNA-seq data (**Extended Data Figure 1)**. Here we demonstrate that our pipeline can effectively use RNA-seq data to reliably generate somatic calls found in whole-exome and whole-genome sequencing. Unlike other existing variant calling pipelines, it does not require a matched normal RNA-seq sample. Many RNA-seq datasets lack matched normal samples because it requires biopsying adjacent tissue and is often difficult to obtain. We also present the additional value of reporting RNA variant allele frequency (VAF) when predicting functional impact of these variants. By comparing the VAFs for variants found in both RNA-seq and WGS, we can further understand the transcriptional mechanisms underpinning these variants and better prioritize driver gene candidates^18^. Our findings indicate that RNA-seq not only complements DNA-based analyses in characterizing expressed somatic variation, but also serves as a valuable resource for researchers to uncover and prioritize potential driver mutations.

## Results

### RNA-seq validates variants detected in WGS

Genomic data from the Pan-Cancer Analysis of Whole Genomes (PCAWG) Consortium^19^ of the International Cancer Genome Consortium (ICGC) and The Cancer Genome Atlas (TCGA) presented a unique opportunity to assess the performance of somatic variant detection using RNA sequencing data. Since PCAWG samples have both whole-genome and RNA-seq data from multiple cancer types, we used this matching data to benchmark existing RNA variant callers by comparing variant calls from RNA-seq data with known somatic variant calls from whole-genome sequencing data. We evaluated the performance of several existing variant callers that can process RNA-seq using simulated and real PCAWG data. We determined that Platypus^20^ had the best speed and performance for variant detection (**Extended Data Figure 2**). We broadly examined the performance of Platypus, alone, for detecting variants in known cancer-associated genes within a specific cancer type. We found that a large number of variants reported by Platypus did not replicate the consensus variants found in the matched WGS data and required a comprehensive filtering strategy to detect true variants (**Extended Data Figure 2**). Starting with Platypus variants, we filtered existing common variants found in dbSNP^21^ and known RNA editing sites from REDIportal^22^. We also generated a panel of normal variants from samples from the Genotype-Tissue Expression^23^ (GTEx) project as another baseline filter (**Table 1**). Further analysis of the candidate variants revealed significant amounts of alignment artifacts, particularly around insertions and deletions and in specific regions of the genome. Gene ontology analysis was performed on potential false positive variants and a striking number were found in immunoglobulin (Ig) and human leukocyte antigen (HLA) genes. Transcripts from these genes often feature extreme levels of diversity, making accurate mapping to these regions difficult for most tools^24,25^. Previous studies also applied similar filters, such as removing variants found at known RNA editing sites and near splice junctions^13^. Other potential false positives were found nearby homopolymer tracts and on reads with multiple variants in close proximity (within 50bp). These events were deemed to be likely sequencing or alignment artifacts and were removed from the candidate variant list.

The above filtering steps were incorporated into our final RNA variant detection pipeline (**Extended Data Figure 1**). We found that the majority of variants found in RNA-seq matched those identified in their whole-genome counterparts and significantly lowered the number of potential false positives. To demonstrate the performance of the pipeline, we show examples in two genes, *NOTCH1* and *NFE2L2*, that have been linked to cancer formation in lung squamous cell carcinoma (LUSC) (**Fig. 1a**). While the variants reported by Platypus, alone, point to a false hotspot mutation, our pipeline largely replicated the WGS mutations found in *NOTCH1*. The *NFE2L2* R34 hotspot mutation was also detected with our pipeline with no other false positives across the rest of the gene.

**Figure 1.**
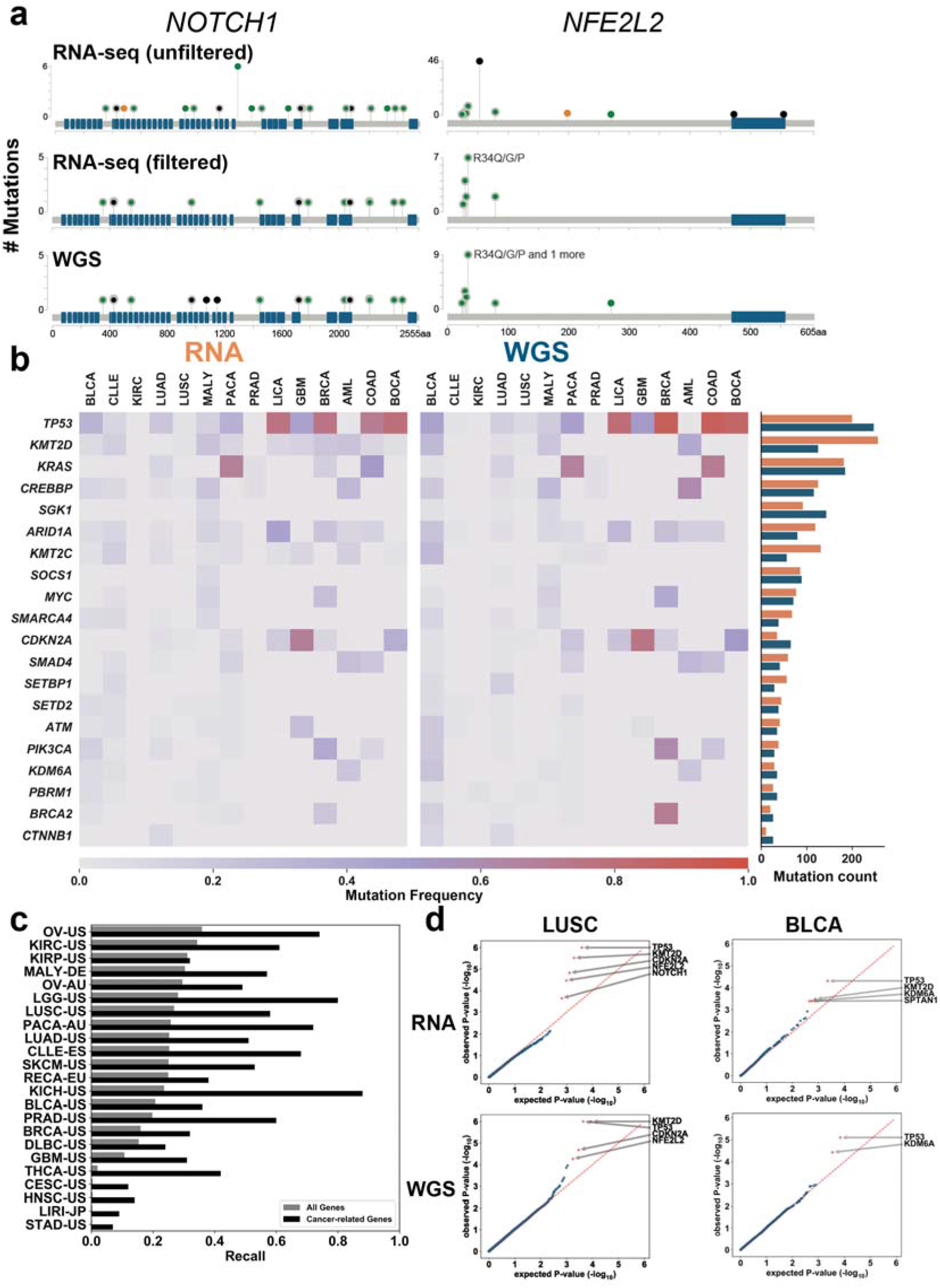
Somatic variants and cancer driver genes identified in RNA-sequencing data. **a.** Stickplot of RNA-seq and WGS mutations found in the genes *NOTCH1* and *NFE2L2* of lung squamous cell carcinoma samples. *NFE2L2* R34* is a known hotspot mutation. **b.** Heatmap compares frequency of samples in a tissue type containing mutations in known cancer-associated genes. Cancer types with large discrepancies in mutation counts were excluded. Barplot displays total number of mutations found in each gene - RNA variants are orange and WGS variants are blue. **c**. Median recall across all tumor types was 0.25. Focusing on a subset of genes provided by the Cancer Gene Census, median recall increases to 0.48. **d**. Quantile-quantile (QQ) plots of p-values generated from oncodriveFML. Red line displays where observed and expected p-values match. Blue dots represent genes with Q-values ≥0.1, red <0.1. Genes with Q-values <0.1 are indicated.

Using this finalized pipeline, we detected 161,809 single-nucleotide somatic variants in all 1,359 RNA-seq samples from the PCAWG dataset (**see Data Availability**). We surveyed several known cancer-associated genes with published hotspot mutations^26^ (*KRAS* G12^27^, *BRAF* V600, *PIK3CA* H1047^28^, etc.) in multiple cancer types to assess the pipeline’s performance in all of the cancer types analyzed by PCAWG. Based on WGS variants found in these genes, we found that 78% of these variant calls were also detected by our pipeline, demonstrating the pipeline’s ability to detect these variants in cancer-related genes while analyzing only RNA-seq data across different tumors. Somatic mutation calls from RNA-seq in *TP53*, the most frequently mutated gene in this study, recapitulated the WGS mutational frequency profile and demonstrated similar high mutational frequencies in liver cancer, colon adenocarcinoma, bone cancer, and breast adenocarcinoma (**Fig. 1b**). Similarly, mutation frequencies and counts in *KRAS*, *MYC*, *CREBBP*, and *SOCS1* were very similar in both RNA-seq and WGS data. Both *KMT2D* and *ARID1A* surprisingly had a larger share of RNA-seq only variants. After individual confirmation with the WGS-aligned reads, the variants were present in both datasets, suggesting that these particular WGS variants were removed during the consensus variant calling process in WGS analysis. Many of these removed variants were flagged as having a strand bias; our pipeline also identifies strand bias, but does not filter these variants due to the existence of stranded RNA-Seq library preparation methods. In lung squamous cell carcinoma (LUSC) samples, our pipeline delivered 7,326 candidate variants; 4,319 of these candidate variants were confirmed in the WGS data. Subsetting for variants within cancer-related genes^29^, revealed 241 candidates shared between the RNA-seq and WGS datasets. Conversely, 177 WGS-derived variants were not detected by our pipeline, demonstrating higher recall within oncogenic loci. This finding was mirrored across all tissue types in the study (**Fig. 1c**).

Previous PCAWG WGS studies identified recurrent noncoding point mutations in multiple genes as being strong candidate drivers^30^. As RNA-seq captures both 5’ and 3’ untranslated regions (UTRs), we decided to test our pipeline’s ability to detect these same UTR mutations. Our pipeline detected somatic variants in the 5’ UTR of *MTG2* and 3’ UTR of *TOB1* and *NFKBIZ* (**Extended Data Figure 3**), but was unable to detect 5’ UTR mutations in *PTDSS1* and *DTL*, as the vast majority of RNA-seq samples had virtually no aligned reads in the specified region of those genes. This may be because somatic variants in this region can often downregulate or upregulate the expression of these genes, particularly in a cancer context^31^. An alternative explanation is that RNA-seq data can exhibit a 3’ end coverage bias due to the cDNA amplification process, resulting in reduced 5’ UTR coverage. Provided there is satisfactory coverage (10 or more reads, **Extended Data Figure 2**), our pipeline is successfully able to detect recurrent UTR variants.

In order to assess our pipeline’s ability to detect cancer driver genes, we used oncodriveFML^32^ to compare the driver mutation profiles of matched RNA-seq and WGS samples in multiple cancer types. OncodriveFML predicts which genes harbor driver mutations using functional impact scores derived from the Combined Annotation Dependent Depletion (CADD) tool. The mean functional impact (FI) score of the mutations within a gene are compared with the distribution of mean functional impact scores of randomly generated mutations. Genes with significant differences in FI scores are likely to be driver genes. We used single nucleotide variants in coding regions from the RNA-seq data across all tumor types to generate driver gene profiles and compared these profiles to their matched WGS samples (**Table 2**). The driver mutation profile of RNA-seq variants called by Platypus alone initially resulted in a multitude of potential driver genes, which can be attributed to the inclusion of germline or false-positive variants. However, RNA-seq variants called from our complete pipeline are often consistent with their WGS equivalents (**Fig. 1d**). Known driver genes were identified such as *TP53*, *KMT2D*, *CDKN2A*, and *NFE2L2* in both lung squamous cell carcinoma (LUSC) RNA-seq and WGS data. In urothelial bladder cancer (BLCA), *TP53* and *KMD6A* were also identified in both RNA-seq and WGS data. The driver gene profiles generated from somatic variants detected by our pipeline largely match the previously reported driver gene profiles from variants found in the corresponding WGS data, demonstrating the ability to use RNA-seq alone to find driver genes.

### RNA-seq identifies allele frequency differences in cancer driver genes

One strength of analyzing variants detected in RNA-seq is the ability to determine the expressed variant allele frequency (VAF) and reveal potential allele bias, providing additional evidence for predicting the functional impact of somatic variants^17,33^. We sought to examine allele bias in the expressed variants from this PCAWG RNA-seq dataset. Somatic variants detected in RNA-seq were selected and profiled for changes in variant allele frequency. We found 161,809 somatic variants in RNA-seq and compared them to 46,607,229 somatic variants in WGS (**see Data Availability**). We saw that the VAFs of the RNA-seq variants also found in WGS had a higher median VAF compared to all RNA variants (**Fig. 2a**). The low VAF RNA variants could also represent variants missed in WGS due to low tumor purity or subclonality as well as sequencing artifacts commonly seen in RNA-seq. The distributions of VAFs higher than 0.5 were similar between all RNA variants and the variants found in both RNA-seq and WGS, confirming our pipeline’s ability to detect somatic variants in highly expressed genes in clonally dominant tumors. *TP53* was the most frequently mutated gene, so we further examined *TP53* variants and found that these variants had higher VAFs overall than either all RNA-seq variants or RNA-seq variants validated with WGS, indicating that *TP53* variants are highly expressed when present (**Fig. 2b**). We also found that *TP53* VAFs can vary among different tumor types. We saw high VAF in BLCA, OV, and STAD, while seeing wider VAF distributions in GBM, HNSC, and LUAD (**Extended Data Figure 4**).

**Figure 2.**
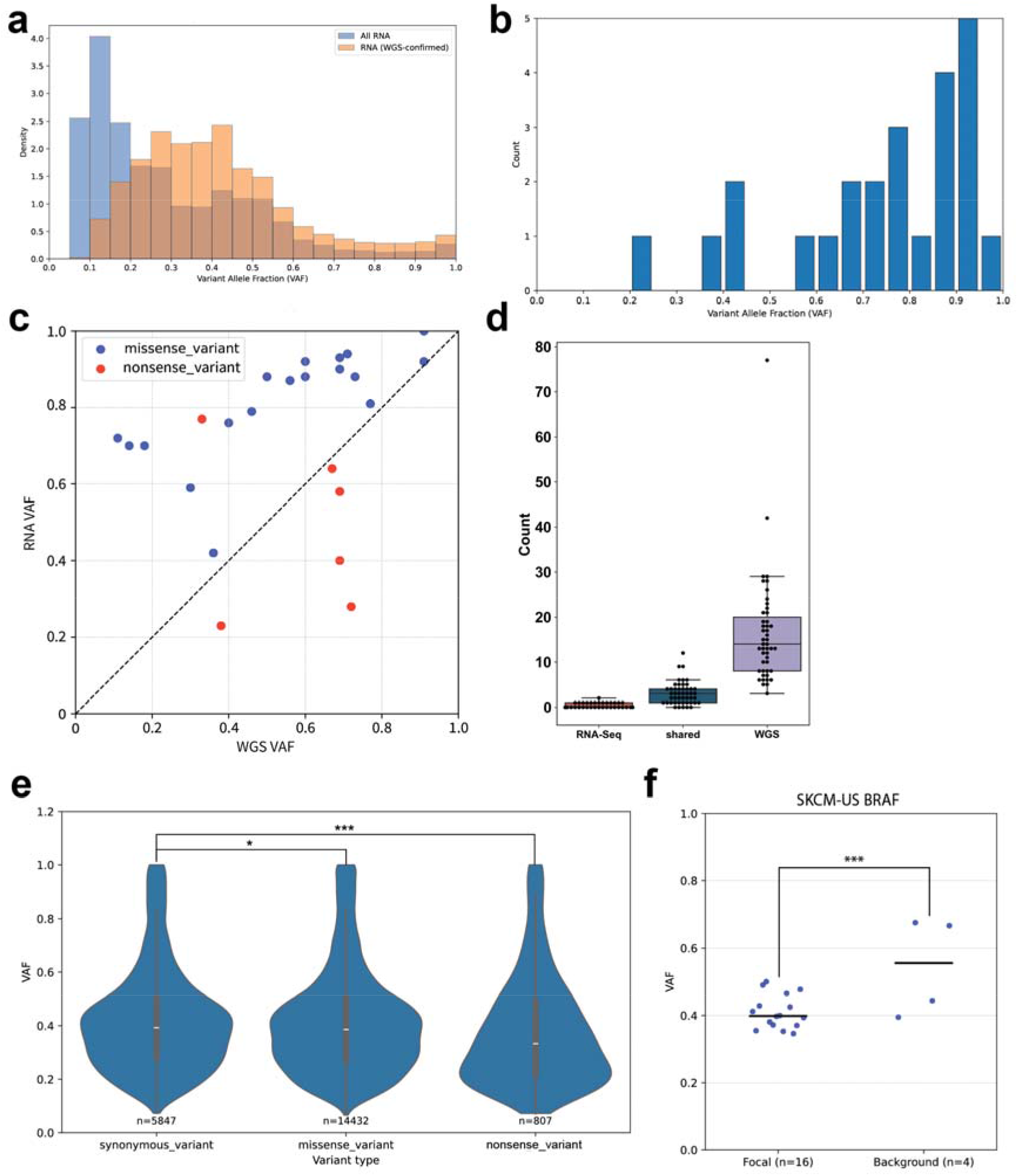
Variant allele frequency in driver genes is influenced by variant impact. a. Distributions comparing VAFs of all variants detected in RNA-seq with VAFs of variants detected in both RNA-seq and WGS. b. Distribution of VAFs of TP53 variants found in breast adenocarcinoma (BRCA). c. Scatterplot of RNA and DNA VAF of TP53 variants found in BRCA. Variants in red are predicted nonsense mutations and variants in blue are predicted missense mutations. D. Boxplot of number of nonsense variants detected in RNA-seq, WGS, or both RNA and WGS data from lung squamous cell carcinoma. e. Violin plot of VAF of variants found in both WGS and RNA-seq, split by synonymous, missense, or nonsense predicted function. VAF distributions between synonymous and missense variants were significantly different with a p-value of 0.0185 by t-test. VAF distributions between synonymous and nonsense variants were also significantly different with a p-value of 2.3719E-08 by t-test. f. Scatterplot of VAF of BRAF variants in skin cutaneous melanoma.

In breast adenocarcinoma (BRCA), we found the majority of *TP53* missense variants (17/18) had an RNA VAF > 0.5, indicating that these variants are likely to have a functional impact **(Fig. 2c)**. We also found that missense variants had higher VAF in RNA compared to DNA while nonsense variants had lower VAF in RNA. When looking at specific TP53 variants, we observed how RNA VAF diverges from the genomic frequency of the alleles (**Extended Data Figure 5**). The combination of RNA and WGS VAF can add further context to the impact of the variant, for example a low VAF WGS variant paired with a high VAF RNA variant could represent an allele-biased mutation found in a low purity tumor.

Consistent with these findings, we were unable to detect most nonsense mutations with RNA data that were previously found in WGS data (e.g. lung squamous cell carcinoma (LUSC), **Fig. 2d**). When we further analyzed which mutation types were not found across the whole RNA-seq dataset, we again found that many missed variants were nonsense mutations, suggesting that they lead to transcript degradation via nonsense-mediated decay^34^, reducing transcript expression and making variants harder to detect through RNA due to low coverage.

While variants in highly expressed oncogenes are confidently detected, the pipeline is unable to detect mutations in potential driver genes with low expression (**Extended Data Figure 6**). In 22 cancer types, we calculated the VAF for 21,086 synonymous, missense, or nonsense mutations that were found both in the RNA-seq and WGS (**Fig. 2e**). We found the VAF for the nonsense variants found in RNA-seq to be significantly lower than both the synonymous and missense variants across all tumor types.

VAFs differ across cancer types, with chronic lymphocytic leukemia (CLLE) having the highest median VAF and prostate adenocarcinoma the lowest (**Extended Data Figure 7**). The VAF distributions for variants found in RNA and WGS were similar across the cancer types; however, for variants only found in RNA, the VAF distributions differed among cancer types. For example, VAFs from variants found only in CLLE RNA-seq samples, had a much lower median VAF compared to variants found in both CLLE RNA-seq and CLLE WGS, which likely indicates that significant number of these RNA-only variants being false positives due to technical artifacts or sequencing issues.

To further examine whether the RNA VAF relates to variant function, we examined whether the same variant in known cancer-related genes have similar RNA VAFs when present in multiple tumor samples. For the 37 recurrent mutations with both WGS and RNA variant calls, we found that their RNA VAFs were less than 0.5 and often clustered together (**Extended Data Figure 8**). Some genes exhibited variability where all samples containing a variant in *BCL2* in non-Hodgkin’s lymphoma (MALY-DE) had a median VAF above 0.5. In non-Hodgkin’s lymphoma, *BCL2* is often highly expressed and high VAF *BCL2* variants are often associated with poor outcomes^35^. In SKCM, we found that the median VAF of *BRAF* V600E variants, a well-known driver mutation, was significantly lower in variants found in both RNA and WGS compared to the VAF of non-canonical (non-hotspot/focal) variants found across the gene (t-test corrected p-val < 0.00489, **Fig. 2f**). Our findings demonstrate somatic variants from RNA can provide additional information when identifying and curating driver mutations across mutation class and tumor type.

## Discussion

Using matching transcriptomic and genomic data from PCAWG Consortium of the ICGC and TCGA projects provided a unique opportunity to evaluate the utility of variant calling RNA-seq data. In this study, we illustrate that somatic variant calling from RNA-seq data, alone, can be used to identify cancer driver genes, identify canonical hotspot mutations, and explore changes in expressed variant allele frequency.

One major challenge in RNA-based variant calling is the high false positive rate associated with alignment artifacts, RNA editing, and coverage biases linked to gene expression. Coverage in RNA-seq is often highly variable and tied to gene expression levels, which limits detection of variants in lowly expressed genes or transcripts found in rare tumor subclones. In our evaluation of existing variant callers, we found that Platypus and other tools report many false positives, requiring a robust filtering strategy to dramatically reduce the number of candidate variants.

After comprehensive filtering for RNA editing sites, common germline variants, and difficult-to-map r egions in immunoglobulin and HLA genes, the majority of somatic variants detected in RNA-seq were also supported by WGS. Our pipeline detected many WGS-derived variants in known cancer genes, including recurrent hotpots in *BRAF*, *KRAS*, *PIK3CA*, and *TP53*. Driver gene analyses using oncodriveFML further demonstrated RNA-seq variants reproduce driver profiles seen in WGS, properly highlighting *KMT2D*, *CDKN2A*, *NFE2L2*, *KDM6A*, and *TP53* as driver genes. These findings validate RNA-seq when variant detection is carefully limited to sufficiently expressed genomic regions.

Beyond variant detection, one key strength of RNA-seq-based variant detection is the ability to measure expressed variant allele frequency (VAF), allowing for identification of allele-specific expression and other post-transcriptional regulatory events. Our pipeline detected many variants with low expressed VAF that were not found in WGS, which may reflect low coverage or technical artifacts. However, they also could represent biologically relevant variants present in rare subclones or low-purity tumors. At higher VAF levels, the number of RNA variants reported was more concordant with WGS variants and indicates that variants in highly expressed genes and clonally dominant tumors were appropriately detected. Variants with high RNA VAF and low DNA VAF may reveal allele specific expression or allelic imbalance. This could point to changes in copy number or some other cis-regulatory advantage, all of which are common biomarkers for tumorigenesis and disease prognosis. Detecting increased expression or copy number gain in conjunction with altered RNA VAF could also further validate oncogenic potential. Similarly, even variants missing from transcriptomic data might help unravel the mechanisms behind a loss of function mutation. Paired with detection of loss of heterozygosity, promoter methylation, or copy number loss, these DNA-only variants would likely be damaging loss-of-function mutations.

Additionally, the RNA-seq variants provide transcript-specific information, as the VAF of these variants reflects their predicted functional impact. We also detected recurrent noncoding mutations in 5’ and 3’ untranslated regions that are typically not captured by WES and identified potential somatic driver mutations that are consistent with previous reports. UTRs are responsible for gene expression regulation and while their connections to disease are still being understood, mutations in these regions have been previously linked to cancer formation^36,37^.

Despite these strengths, several limitations warrant consideration. RNA-seq–based variant detection is inherently limited to expressed regions and is sensitive to gene expression level, transcript isoform usage, and sequencing depth. Variants in lowly expressed genes are less likely to be detected. Variants associated with transcript degradation, particularly nonsense mutations, are also hard to detect. RNA VAF also does not directly reflect clonal prevalence, as it is influenced by transcriptional regulation, allele-specific expression, and RNA stability. Alignment challenges remain in repetitive, polymorphic, or highly homologous genomic regions, even after careful variant filtering.

Future work should focus on integrating RNA-seq–derived variant information with additional molecular layers, including copy number, methylation, and single-cell expression data, to better disentangle transcriptional effects from underlying genomic architecture. Joint modeling of RNA VAF with DNA VAF, copy number state, and gene expression may improve inference of allele-specific regulation and functional impact. Extending this framework to single-cell and spatial transcriptomic data could further resolve the cellular contexts in which somatic variants are expressed. Finally, incorporating RNA-derived VAF into driver mutation prioritization frameworks may improve identification of functionally relevant variants, particularly in settings where DNA sequencing is limited.

In summary, our results show that RNA sequencing, when analyzed using a carefully designed and stringent filtering strategy, can recover a substantial fraction of cancer-relevant somatic variants identified by whole-genome sequencing. Beyond variant detection, RNA-derived allele frequency measurements provide complementary information about variant expression and post-transcriptional regulation that is not directly accessible from DNA data alone. Our findings support the use of RNA sequencing as a complementary approach for studying expressed somatic variation and prioritizing functionally relevant mutations, particularly when matched DNA sequencing is unavailable.

## Methods

### Aligned reads processing

We used data from The Pan-Cancer Analysis of Whole Genomes (PCAWG) Consortium of the International Cancer Genome Consortium (ICGC) and The Cancer Gene Atlas (TCGA). We downloaded 1,359 RNA-seq samples from the PCAWG data portal. This dataset features 30 cancer types. 161 of these samples originate from normal solid tissue or tissue adjacent to the tumor. These reads were aligned with STAR^38^ (v2.4.0i) and hs37d5^39^ or GENCODE (release 19)^40^ as the reference gene annotation. Matched WGS data for these samples were used to evaluate pipeline performance. To generate a simulated dataset, 300 randomly selected somatic single-nucleotide variants (SNVs) were manually added to aligned reads from 20 PCAWG normal tissue RNA-seq samples to simulate tumor RNA-seq data. All variants were located in the coding regions of the genome and had random allele frequencies. A test dataset of 8 donors with matched tumor and normal RNA-seq from 8 tumor types were curated and used to evaluate the performance of the variant calling tools.

RNA-seq data from normal tissue samples were downloaded from the Genotype-Tissue Expression (GTEx) project portal and re-aligned with identical parameters. Variants detected in these samples were then used to generate a panel of normal variants. 20 samples from 11 different tissue types were incorporated into this panel (**Table 1**).

### Initial variant caller evaluation

Four open-source variant callers previously reported to be compatible with RNA-seq data were evaluated – Platypus^20^ (v0.8.1.1), GATK^5^ (v4.1.9), VarDict^41^ (v1.5.5), and FreeBayes^42^ (v1.1). We ran the tools using default recommended options and recommended preprocessing steps for Platypus (Opossum^20^) and GATK (SplitNCigarReads). We measured the speed, recall, PPV, and resource requirements of the four variant callers processing 10 RNA-seq samples, comparing the results between the 5 pairs of normal and tumor samples.

### Variant calling pipeline description

To call variants using only RNA-seq data, we created a pipeline we call RNA-VACAY, which is a modular workflow built on Python 2.7 that automates task assignment, downloading and preprocessing data, tool execution, and variant analysis. Each task is completed within a Docker container.

#### 1. Data retrieval

This pipeline is built to specifically handle RNA-seq reads aligned with STAR (**<u>Supplementary Note 1</u>**) currently stored in either PCAWG or TCGA repositories. The download module includes the recommended tools by each consortium. RNA-VACAY accepts file manifests generated by these data repository portals and automates downloading. It can also accept user-generated file manifests to call variants in either previously downloaded data or RNA-seq data generated by the user.

#### 2. Preprocessing data

Aligned reads are sorted and indexed by samtools if necessary and then preprocessed with Opossum (v0.2). Opossum prepares RNA-seq data for variant calling by Platypus, GATK, and other callers. It splits reads mapped across splice junctions and ensures that minimal information is lost at read ends by merging overlapping reads and modifying base qualities at the edges of these reads. Opossum also eliminates duplicate reads.

#### 3. Variant calling

Platypus is a Bayesian haplotype-based variant caller that uses local *de novo* assembly and realigns sequences to detect variants. Platypus also shares variant information between multiple samples, increasing the confidence of calls that are weakly supported in one sample, but strongly supported in related samples.

#### 4. Filtering

Raw variant calls from Platypus are first filtered with a custom panel of normal variants generated from RNA-seq samples from the GTEx repositories. Subsequent filters incorporate a combination of preexisting common and normal variant databases - dbSNP, gnomAD, and REDIportal (RNA editing sites). Variants with low quality scores or sequencing depth (<7) were filtered. Variants found in certain locations, such as known decoy regions, and repeat regions were excluded. Variants found in human leukocyte antigen genes, immunoglobulin genes, and pseudogenes were also excluded. Variants found within 50 bases of other variants with similar allele frequencies or within 10 bases adjacent to homopolymer tracts of 5+ bases were also excluded. An optional filter will prevent removal of variants found in known cancer hotspots, regardless of call quality.

Normal variants from matched normal RNA-seq samples, if available, can also be incorporated as an optional filter.

#### 5. Annotation and analysis

Filtered variants were annotated with SnpEff (v4.3t). SnpEff categorizes the variants based on their genomic locations and predicts the coding effects of these variants.

These candidate variants were then analyzed using custom Python scripts. Driver analysis was performed by oncodriveFML. OncodriveFML calculates a profile of somatic mutations in specific genomic regions and identifies genes that have a higher mutational frequency compared to their background mutation rate. All calls outside of the coding region and any non-single nucleotide variants were filtered before running oncodriveFML.

### PCAWG RNA-seq data analysis

RNA-seq samples were downloaded as cohorts based on cancer type. RNA-VACAY was run on multiple OpenStack instances in parallel. Custom python scripts were developed to handle and aggregate results.

### RNA and DNA variant comparison

The variants in particular genes were visualized as stickplots with cBioPortal^43,44^. Known cancer-related genes in specific cancer types were chosen and plotted using the MutationMapper tool^43,44^. Custom python scripts were written to analyze the overlap between the RNA and WGS variant sets and to calculate the mutational frequencies of the 25 most frequently mutated cancer-related genes across each PCAWG tumor type.

### Driver mutation profiling

OncodriveFML^32^ (v2.2.0) was used to identify genes with potential driver mutations. We ran oncodriveFML with default settings on filtered variants, using the whole-exome sequencing option.

### Allele frequency analysis

RNA-seq and WGS variant calls were merged into unified datasets. Each RNA-seq–derived variant was annotated to determine whether it (1) shared an identical genomic position with a corresponding WGS variant from the same donor and/or (2) overlapped a significant driver mutation identified by OncodriveFML. Variant allele frequency (VAF) was calculated as the proportion of reads supporting the variant allele relative to the total read depth at that locus. Variants detected in both RNA-seq and WGS were further annotated. The functional impact of missense variants was evaluated using PolyPhen-2^45^. Genes harboring overlapping variants in three or more donors within a single tumor type were identified, and their VAF distributions were visualized. Recurrently mutated loci shared across multiple donors within a tumor type were also identified, and the VAFs of these variants were compared with those of other variants within the same gene.

### Splice junction analysis

Filtered variants were annotated with their distance to the nearest splice junction, calculated using exon boundary coordinates from GENCODE v19 (GTF). Variants located within 5 bp of a splice junction were excluded to minimize alignment ambiguity. The remaining variants were cross-referenced with an actionable cancer gene list obtained from Sayles et al., Cancer Discovery^46^ (2019).

### 5’ and 3’ UTR mutation confirmation and visualization

Custom python scripts were written to uncover variants detected by RNA-VACAY that matched previously published genes with recurrent 5’ and 3’ UTR mutations. We used Integrated Genomics Viewer^47^ (v2.8.3) to visually confirm the variants.

## Code Availability

Code for the variant calling and driver variant filtering pipeline is available at https://github.com/BrooksLabUCSC/RNA-VaCay

## Data Availability

All data used in this study was obtained from TCGA (https://portal.gdc.cancer.gov/) and ICGC (https://docs.icgc-argo.org/docs/data-access/icgc-25k-data). RNA variant calls will be deposited in dbGAP.

## Acknowledgements

We thank Cameron Soulette for his valuable support during the genesis of this project and the members of the Brooks and Vaske Lab for their helpful feedback. We also thank Yoseph Barash for helpful suggestions. This work was supported by the University of California Cancer Research Coordinating Committee CRN-19-586125 (A.N.B.). J.A. was partially supported by an NIH training grant 5T32HG008345 and the UCSC STARS Re-entry Scholarship. J.L.A. was funded by the University of Leeds (University Academic Fellow scheme). Partial support was also provided by the following awards: NIH/NCI 1R21CA238600-01(A.N.B.) and Pew Charitable Trusts (A.N.B.).

## Author Contributions

Jon Akutagawa: Conceptualization, Visualization, Data Curation, Investigation, Formal analysis, Software, Methodology, Writing - Original Draft

Allysia J Mak: Visualization, Data Curation, Investigation, Formal analysis, Software

Tanvi Damle: Visualization, Data Curation, Investigation, Formal analysis, Validation, Writing - Review & Editing

Colette Felton: Conceptualization, Supervision, Writing - Original Draft, Writing - Review & Editing

Julie L Aspden: Conceptualization, Supervision

Angela N Brooks: Conceptualization, Supervision, Project administration, Funding Acquisition, Writing - Review & Editing

## Supplementary Tables

**Table 1** - List of GTEX samples used for panel of normals. List of samples selected from GTEx that represent the panel of normals generated for filtering somatic variants.

**Table 2** - List of oncodriveFML genes for each tumor type (only significant genes). Genes containing driver mutations predicted by oncodriveFML.

---

**Extended Data Fig. 1.**
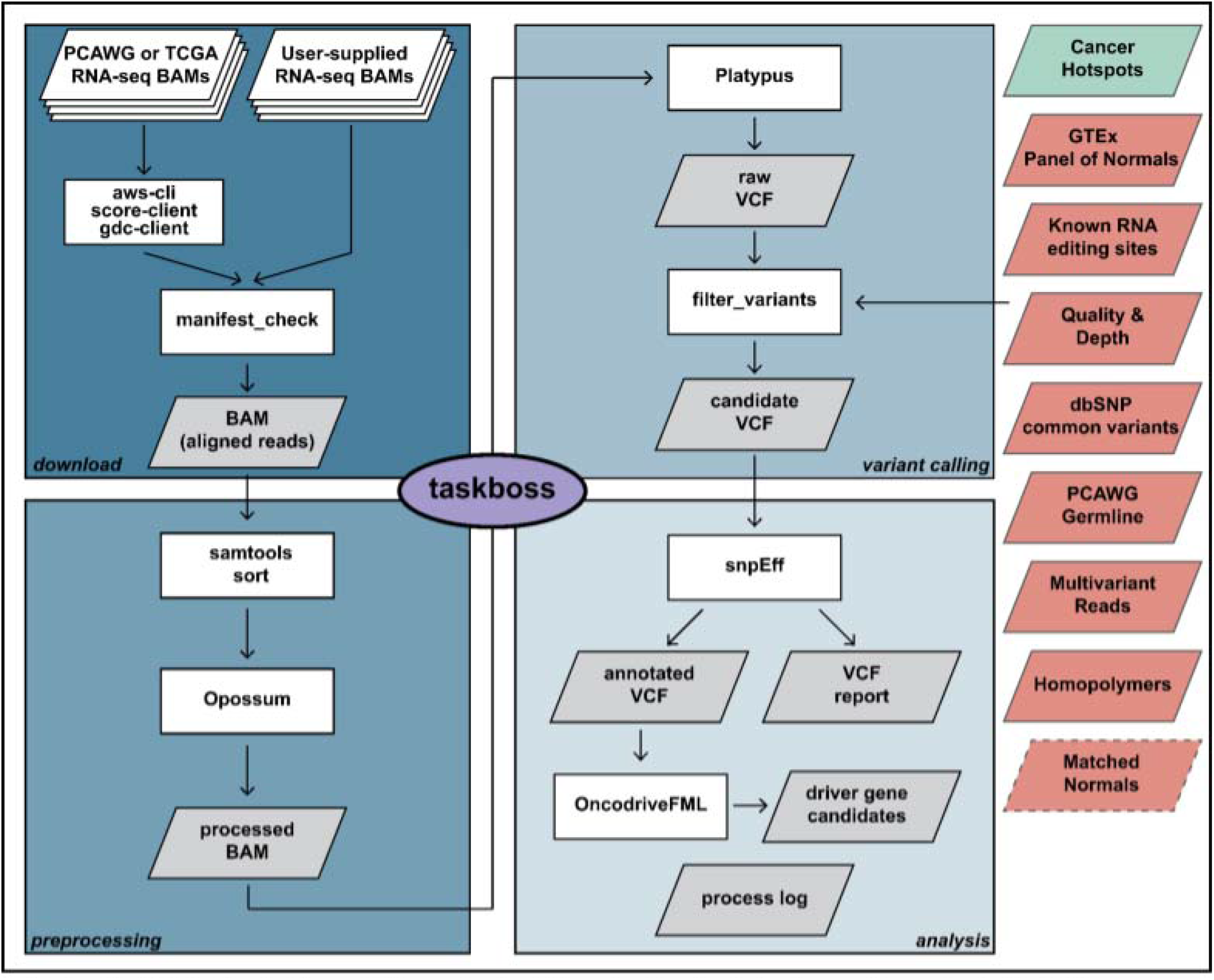
A scalable variant calling pipeline to detect somatic variants in RNA-seq data. Schematic of a somatic variant calling pipeline designed for RNA-seq data. The pipeline consists of 4 main modules - data download, preprocessing, variant calling, and analysis. Taskboss, a controller module, assigns tasks according to available resources and can assist in resuming pipeline operations after interruptions. Multiple filtering options (green and red) can be toggled to keep known cancer mutations and remove likely false positives. Variants from matched normal samples, if available, can also be used to filter common variants. White boxes refer to components of the pipeline and gray parallelograms refer to outputs generated by the pipeline.

**Extended Data Fig. 2.**
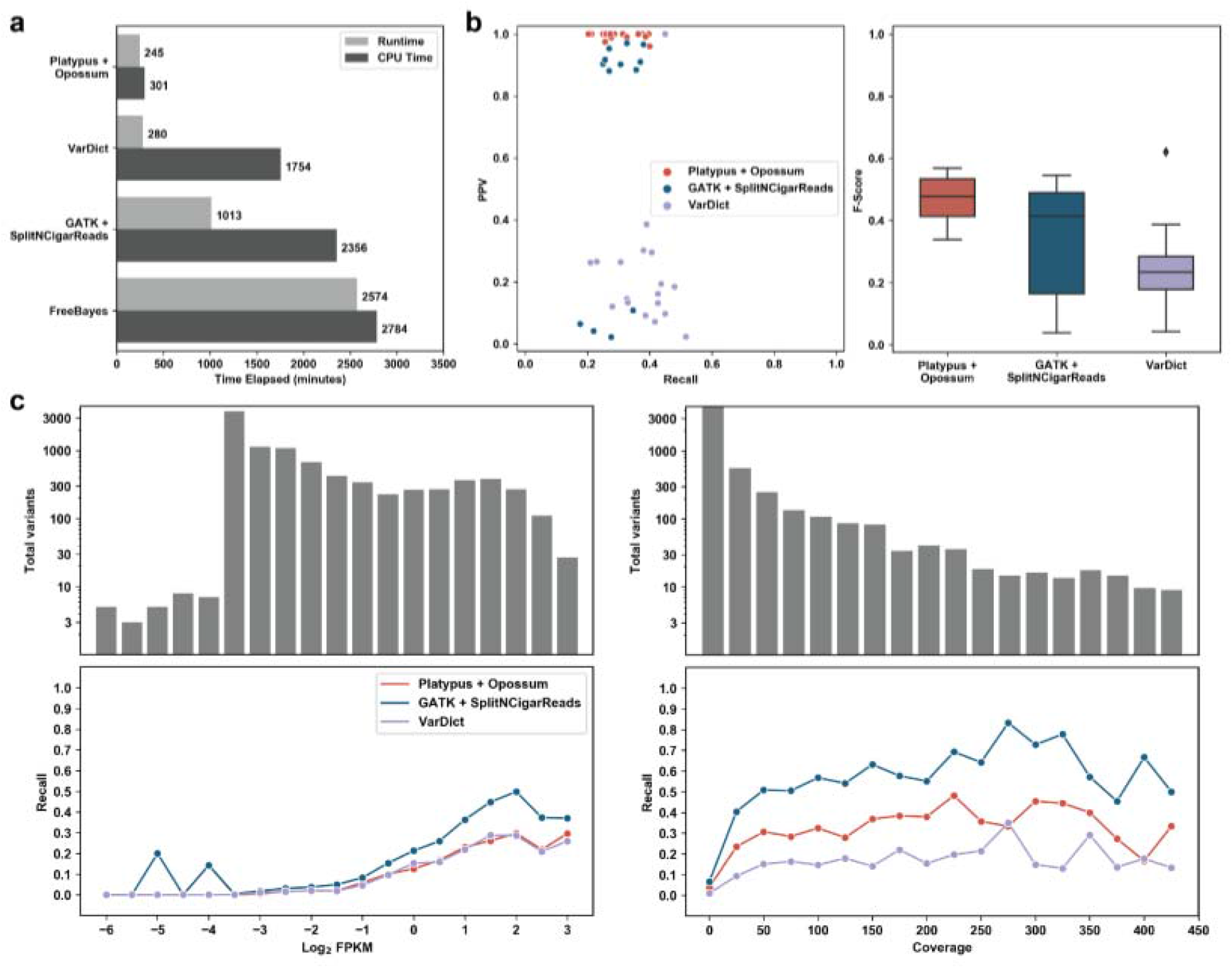
Platypus is the tool best fit for a fast and sensitive variant calling pipeline. **a.** Four variant callers were evaluated for runtime using 5 synthetic RNA-seq samples. The mean time was reported. Together Platypus and Opossum - a preprocessing step - were significantly faster and consumed less resources than all other variant callers. **b.** Three variant callers were evaluated for recall and positive predictive value (PPV) with a synthetic RNA-seq dataset of aligned reads with spiked-in somatic mutations. Platypus had the highest median F-score. **c.** The three tools were then evaluated with a small test dataset (12 samples from diverse cancer types) curated from PCAWG. All variant callers generally were more sensitive in genes with higher expression (log_2_ FPKM) and variants with higher coverage.

**Extended Data Fig. 3.**
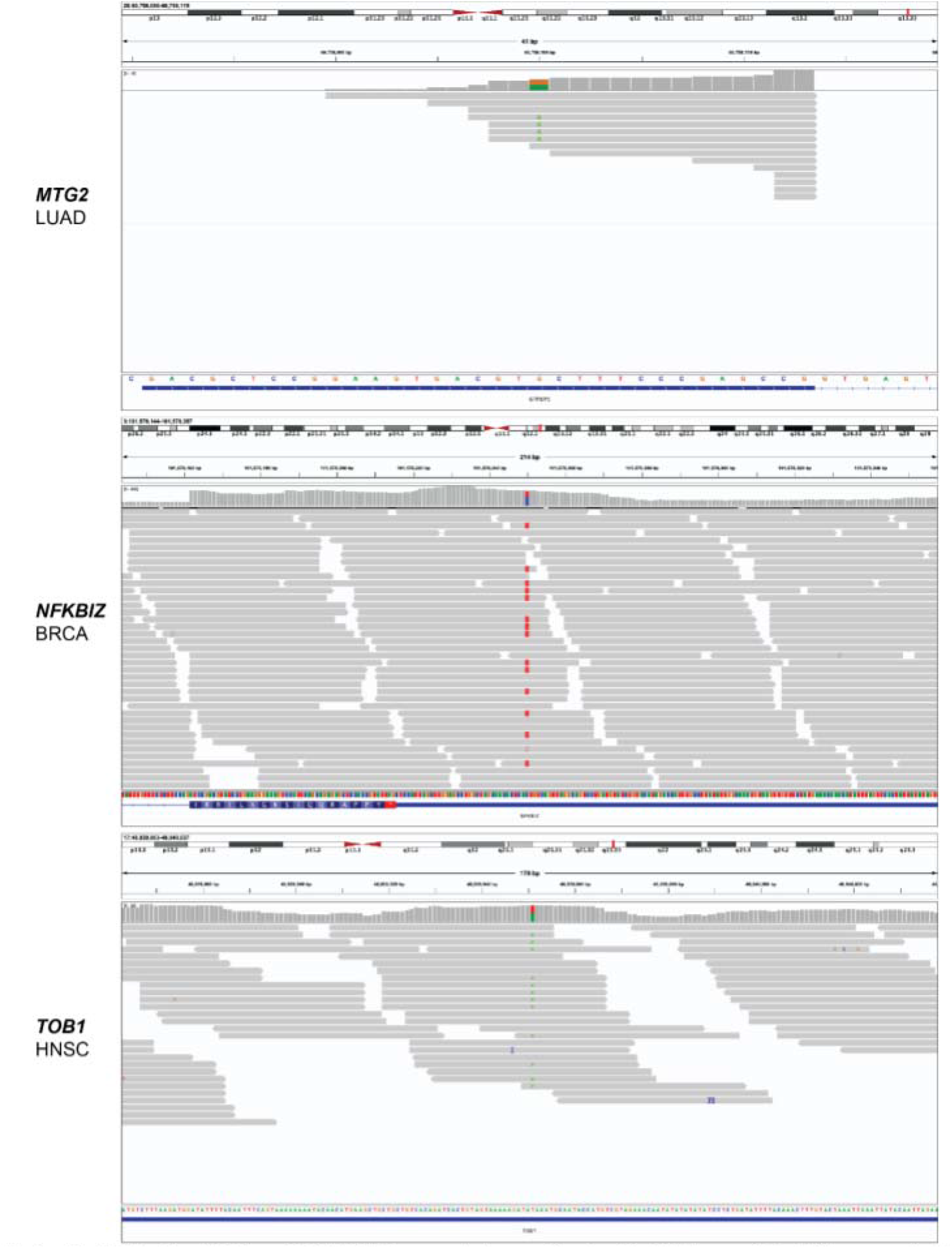
5’ and 3’ UTR driver mutations detected in RNA-seq data. IGV screenshots confirm detection of previously reported candidate driver mutations in the 5’ UTR of *MTG2* and and 3’ UTR of *TOB1* and *NFKBIZ*.

**Extended Data Fig. 4.**
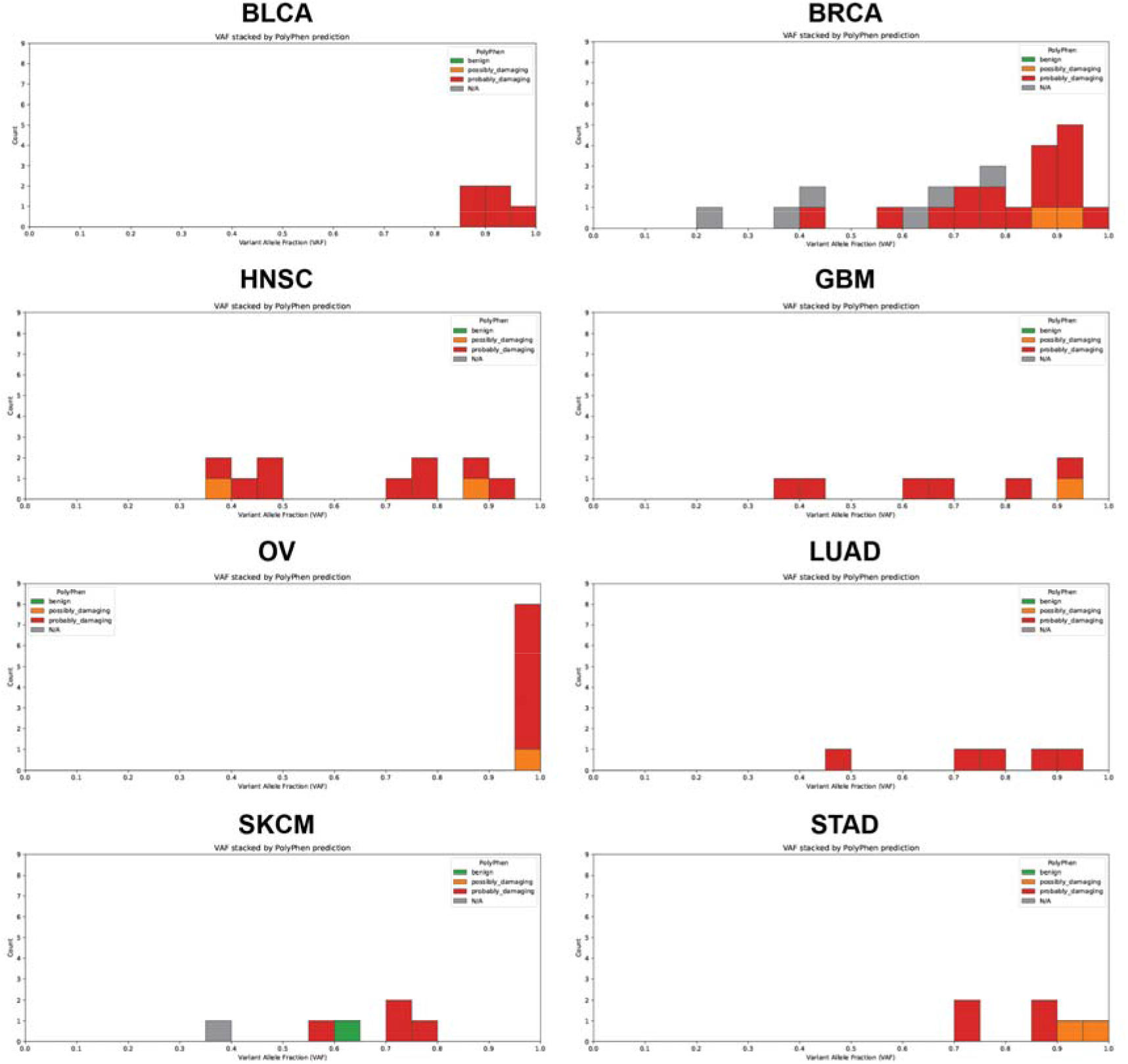
VAF of TP53 variants vary across cancer types. VAF distribution histograms of TP53 variants in bladder adenocarcinoma, breast adenocarcinoma, head and neck squamous cell carcinoma, glioblastoma, ovarian cancer, lung adenocarcinoma, skin cutaneous melanoma, and stomach adenocarcinoma. Histograms are colored by Polyphen2 predicted functional impact.

**Extended Data Fig. 5.**
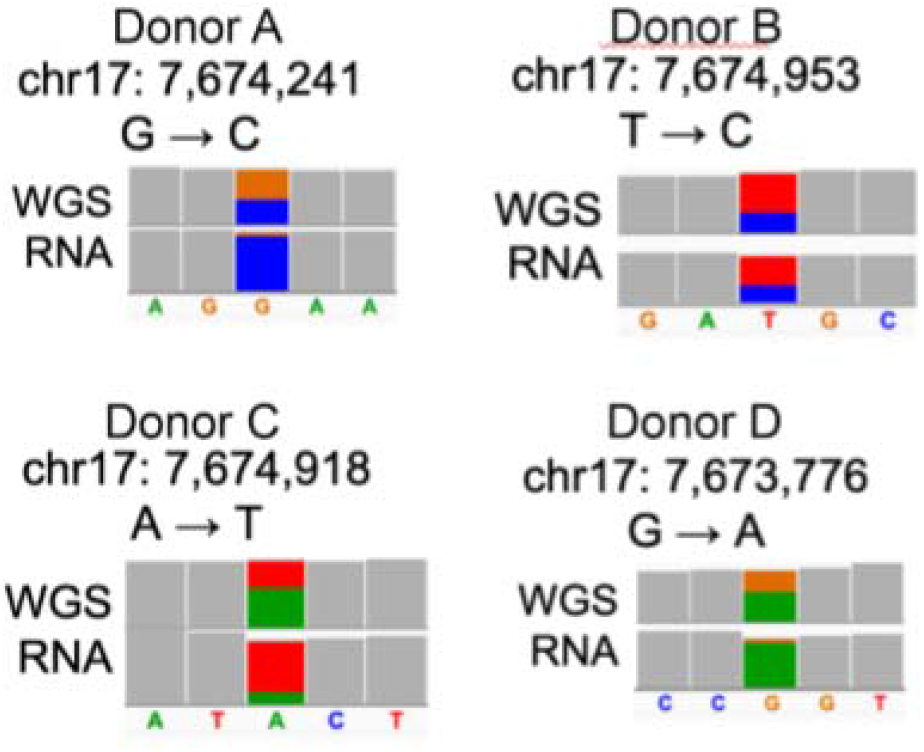
RNA VAF of *TP53* variants diverges from genomic VAF. IGV figures displaying aligned read counts for *TP53* variants found in WGS and RNA-seq data.

**Extended Data Fig. 6.**
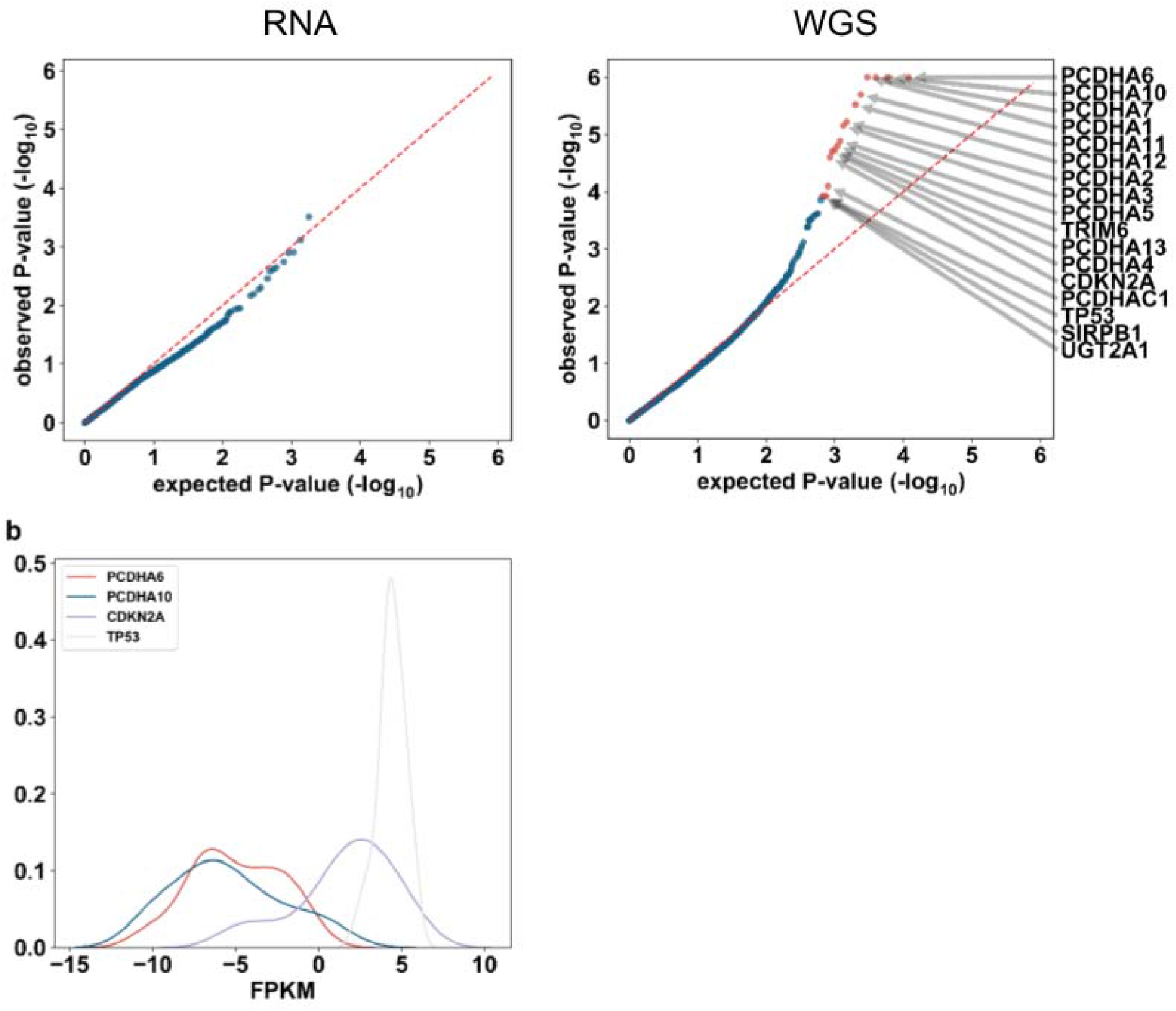
Potential driver genes contain variants found in lowly expressed genes. **a.** Quantile-quantile (QQ) plots of p-values generated from oncodriveFML in skin cutaneous melanoma (SKCM). **b.** Density plot of gene expression of top WGS driver gene candidates in SKCM samples.

**Extended Data Fig. 7.**
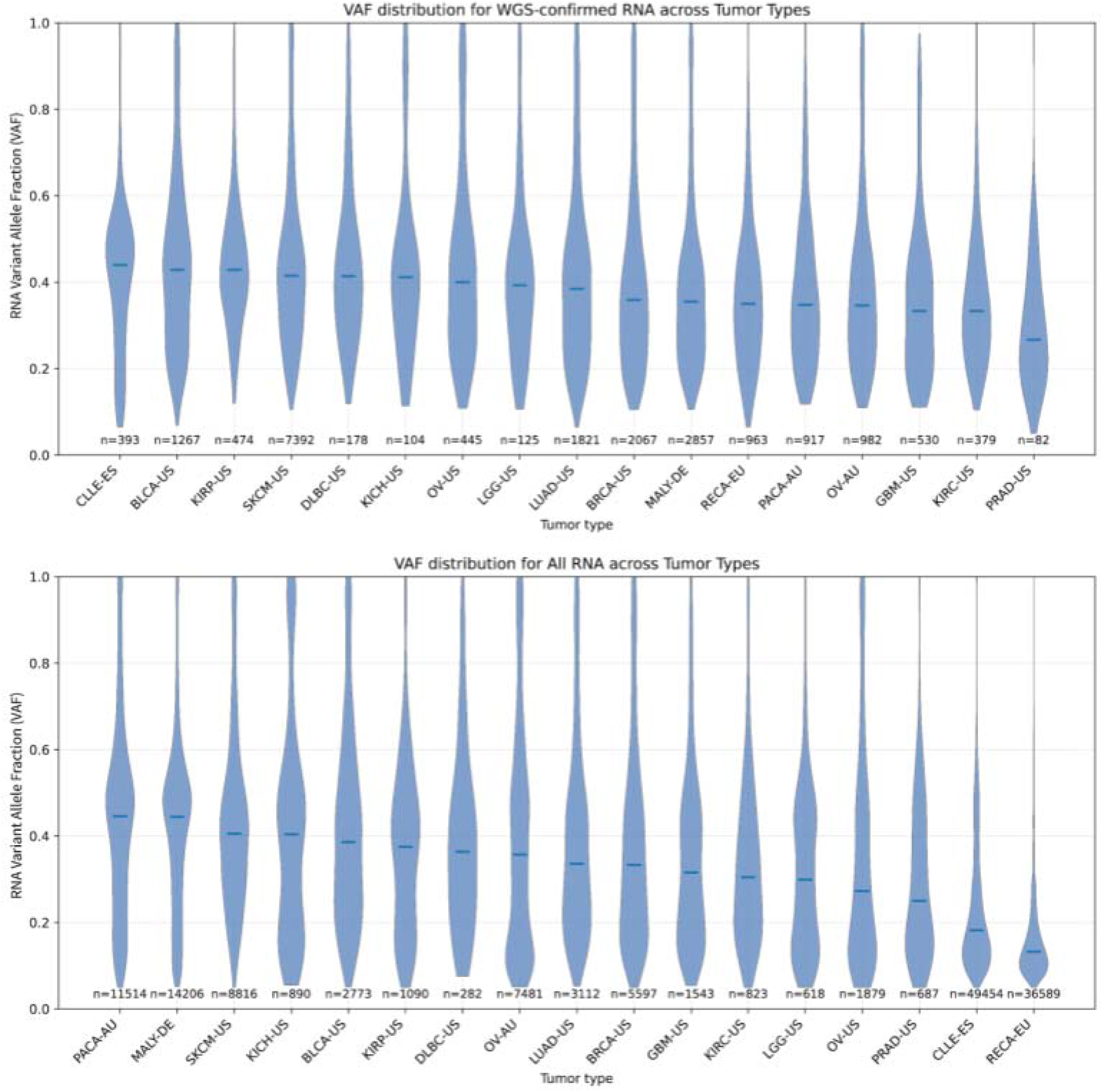
Variant allele frequencies of somatic variants detected in both RNA and WGS have similar distributions across cancer types. Violinplots of RNA VAFs of variants detected in both RNA-seq and WGS (top) and detected only in RNA-seq (bottom).

**Extended Data Fig. 8.**
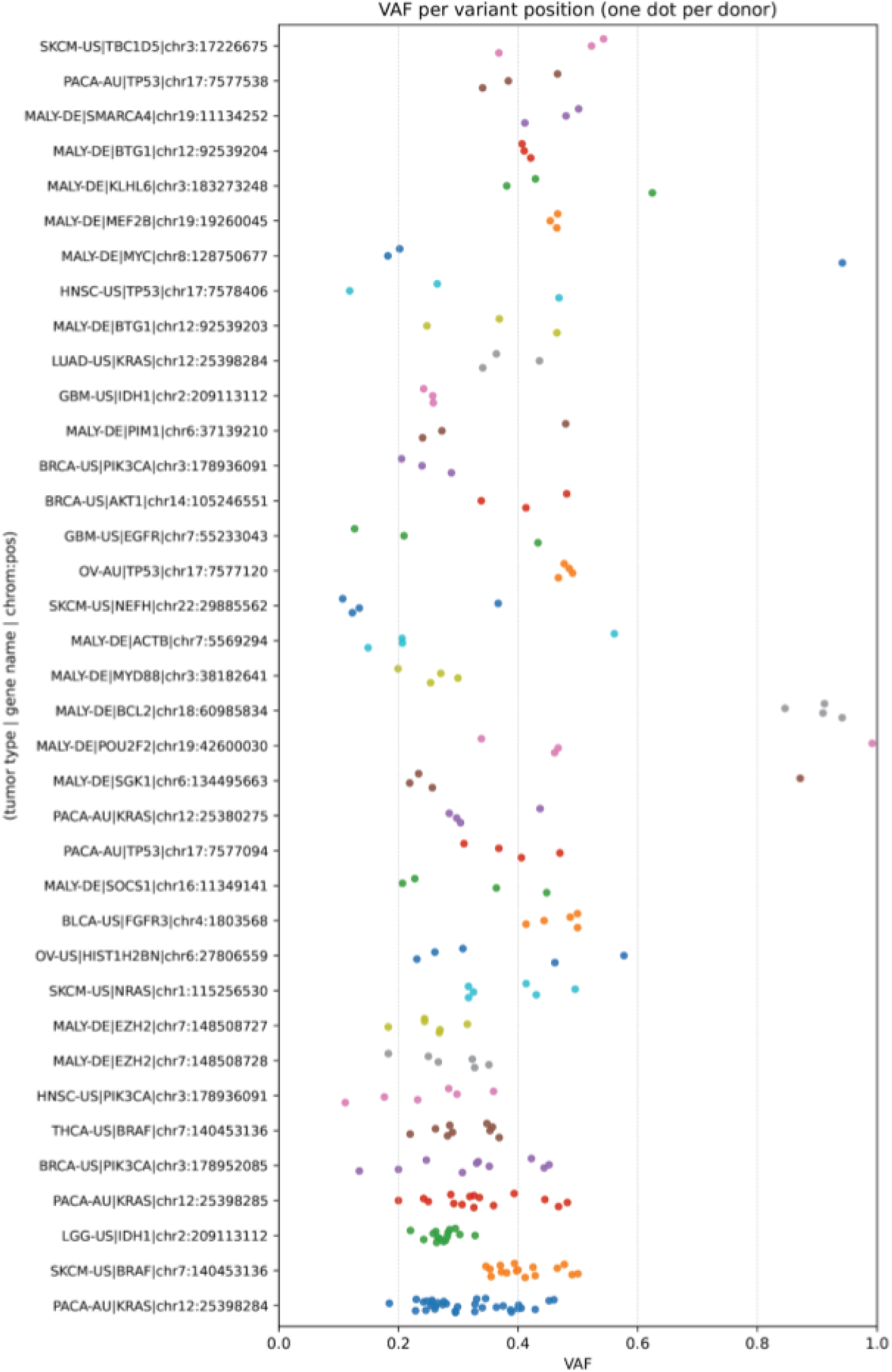
VAF of cancer-related gene variants vary across cancer types. Scatterplot of VAF of variants detected in known cancer-related genes.

