## Supplementary Information for "RNA-sequencing as an expression-informed alternative for somatic variant detection and cancer driver identification"

### Supplementary Note 1. Sample STAR command

~/STAR --genomeDir ~/reference_indices/hs37/star_genome \

--readFilesIn ~/sample_1.fastq ~/sample_2.fastq \

--runThreadN 8 \

--outFilterMultimapScoreRange 1 \

--outFilterMultimapNmax 20 \

--outFilterMismatchNmax 10 \

--alignIntronMax 500000 \

--alignMatesGapMax 1000000 \

--sjdbScore 2 \

--alignSJDBoverhangMin 1 \

--genomeLoad NoSharedMemory \

--limitBAMsortRAM 0 \

--outFilterMatchNminOverLread 0.33 \

--outFilterScoreMinOverLread 0.33 \

--sjdbOverhang 100 \

--outSAMstrandField intronMotif \

--outSAMattributes NH HI NM MD AS XS \

--outSAMunmapped Within \

--outSAMtype BAM SortedByCoordinate \

--outSAMheaderHD @HD VN:1.4 \

--twopassMode Basic \

--twopass1readsN -1 \

> ~/star_output.log

### Supplementary Note 2. Reference files used in RNA-VACAY

**repeat_masker_hg19.no_chr.bed** – a BED file generated from the RepeatMasker track found in the UCSC Genome Browser

**pon.no_header.vcf** – a VCF of variants found in GTEx samples (not included)

**00-common_all.vcf.gz** – a VCF of common variants from the dbSNP database (hg19)

**pcawg8.snps.indels.svs.phased.tcga.v2.controlled.vcf.gz** – a VCF of germline variants found in TCGA samples (not included)

**pcawg8.snps.indels.svs.phased.icgc.v2.controlled.vcf** – a VCF of germline variants found in ICGC samples (not included)

**pcawg_normal.vcf** – a VCF of variants found in normal PCAWG samples (not included)

**publication_hotspots.vcf** – a VCF of known cancer hotspots (hg19)

**TABLE1_hg19.txt** – a TXT with a list of known RNA editing sites from REDIportal

**02_cds.regions.bed** – a BED file with CDS regions (hg19)

### Supplementary Note 3. Sample RNA-VACAY commands

**PCAWG manifests and metadata**:

python rna-vacay.py -pc ~/manifests/sample_manifest.tsv -pm ~/metadata/sample_metadata.tsv > rna_vacay_output.log

**User-generated or previously downloaded aligned RNA-seq reads**:

python rna-vacay.py -bm ~/manifests/sample_manifest.tsv > rna_vacay_output.log
