## Supplementary Table 1 for "RNA-sequencing as an expression-informed alternative for somatic variant detection and cancer driver identification"

### Supplementary Table 1. GTEx samples used to generate the panel of normals.

| **Biospecimen Repository Sample ID** | **Histological Type** |
| --- | --- |
| GTEX-XV7Q-2626-SM-4BRVA | Adipose Tissue |
| GTEX-XBED-2226-SM-47JYQ | Skin |
| GTEX-XV7Q-1426-SM-4BRWA | Ovary |
| GTEX-XYKS-1726-SM-4E3IO | Ovary |
| GTEX-XQ3S-0526-SM-4BOQA | Heart |
| GTEX-VUSG-2626-SM-4KKZI | Muscle |
| GTEX-T5JW-2026-SM-4DM63 | Breast |
| GTEX-T5JC-1526-SM-4DM68 | Kidney |
| GTEX-13CF3-2126-SM-5IFJP | Breast |
| GTEX-11LCK-1926-SM-5A5KE | Small Intestine |
| GTEX-1339X-0626-SM-5IJER | Lung |
| GTEX-Q734-0526-SM-2I3EH | Thyroid |
| GTEX-P4PQ-0526-SM-2HMKR | Lung |
| GTEX-OOBK-1626-SM-2HMKG | Muscle |
| GTEX-OXRK-0926-SM-2HMKP | Lung |
| GTEX-NPJ8-0011-R4a-SM-2HML3 | Brain |
| GTEX-N7MS-0011-R7a-SM-2HMKN | Brain |
| GTEX-QDVN-2126-SM-2I3FR | Adipose Tissue |
| GTEX-OHPM-0326-SM-2HMKT | Heart |
| GTEX-Q2AG-0126-SM-2HMLB | Skin |
